# Effects of prey turnover on poison frog toxins: a landscape ecology approach to assess how biotic interactions affect species phenotypes

**DOI:** 10.1101/695171

**Authors:** Ivan Prates, Andrea Paz, Jason L. Brown, Ana C. Carnaval

## Abstract

Ecological studies of species pairs demonstrated that biotic interactions promote phenotypic change and eco-evolutionary feedbacks. However, we have a limited understanding of how phenotypes respond to interactions with multiple taxa. We investigate how interactions with a network of prey species contribute to spatially structured variation in the skin toxins of the Neotropical poison frog *Oophaga pumilio*. Specifically, we assess how beta-diversity of alkaloid-bearing arthropod prey assemblages (68 ant species) and evolutionary divergence among populations (from a neutral genetic marker) contribute to frog poison dissimilarity (toxin profiles composed of 230 different lipophilic alkaloids sampled from 934 frogs at 46 sites). We show that ant assemblage turnover predicts alkaloid turnover and unique toxin combinations across the range of *O. pumilio*. By contrast, evolutionary relatedness is barely correlated with toxin variation. We discuss how the analytical framework proposed here can be extended to other multi-trophic systems, coevolutionary mosaics, microbial assemblages, and ecosystem services.

## Introduction

Phenotypic variation within species, an essential component of evolutionary theory, has received increased attention by ecologists (Bolnick *et al*., 2011; Vildenes and Langangen, 2015). This interest has been chiefly motivated by evidence showing that phenotypic change, both adaptive and plastic, can happen within contemporary time scales and thus has consequences for ecological processes (Shoener, 2011; Hendry, 2015). Changes in trait frequencies can affect survival and reproduction and ultimately determine population density and persistence of a given species. In turn, these demographic changes may influence community-level and ecosystem functions such as nutrient cycling, decomposition, and primary productivity (Miner *et al*., 2005; Pelletier *et al*., 2009; Post and Palkovack, 2009). This interplay between evolutionary and ecological processes, or ‘eco-evolutionary dynamics’, has brought phenotypes to the center of ecological research (Hendry, 2015).

Several studies focusing on interacting species pairs have demonstrated that population-level phenotypic change can originate from biotic interactions, leading to geographic trait variation within species. For instance, different densities of Killifish predators in streams lead to distinct morphological and life history traits in their Trinidadian guppy prey (Endler, 1995). In western North America, levels of tetrodotoxin resistance in garter snake predators can match toxicity levels in local populations of tetrodotoxin-defended newt prey (Brodie *et al*., 2002). By focusing on the associations between two species, these studies have provided crucial insights into how interactions can lead to trait divergence and potentially shift the evolutionary trajectory of natural populations (Post and Palkovack, 2009; Hendry, 2015). The resulting trait diversity can have broad ecological consequences, altering the role of species in the ecosystem at a local scale (Palkovacs *et al*., 2009).

However, we still have a limited understanding of how interactions among multiple co-distributed organisms contribute to complex phenotypes, particularly when the set of interacting species and their phenotypes vary geographically. An example of a complex phenotype that is shaped by a network of interactions with many species is the chemical defense system of poison frogs (Dendrobatidae). In this clade of Neotropical amphibians, species can exhibit dozens to hundreds of distinct lipophilic alkaloid toxins in their skin, and aposematic color patterns advertise their distastefulness (Saporito *et al*., 2012; Santos *et al*., 2016). Poison composition varies spatially within species such that populations closer in geography tend to have alkaloid profiles more similar to one another than to populations farther away (Saporito *et al*., 2006, 2007a; Mebs *et al*., 2008; Stuckert *et al*., 2014). This geographic variation in toxin profiles has been attributed to local differences in prey availability, because poison frogs obtain their defensive alkaloids from dietary arthropods (Daly *et al*., 2000; Saporito *et al*., 2004, 2007b, 2009; Jones *et al*., 2012). For instance, specific toxins in the skin of a frog may match those of the arthropods sampled from its gut (McGugan *et al*., 2016). However, individual alkaloids may be locally present in frog skins but absent from the arthropods they consume, and vice-versa; thus, the extent to which frog chemical traits reflect arthropods assemblages remains unclear (Daly *et al*., 2000, 2002; Saporito *et al*., 2007b; Jones *et al*., 2012). Population differences may also stem from an effect of shared evolutionary history, because alkaloid sequestration may be partially under genetic control in poison frogs (Daly *et al*., 2003, 2005, 2009). These drivers of toxin turnover can have broad consequences for the ecology of poison frogs, because alkaloids protect these amphibians from numerous predators (Darst and Cummings, 2006; Gray *et al*., 2010; Weldon *et al*., 2013; Murray *et al*., 2016), ectoparasites (Weldon *et al*., 2006), and pathogenic microorganisms (Macfoy *et al*., 2005).

To address the question of how interactions among multiple organisms contribute to spatially structured phenotypes, we investigate how prey assemblage turnover and evolutionary divergence among populations predict the rich spectrum of toxins secreted by poison frogs. We focus on the well-studied toxin profiles of the strawberry poison frog, *Oophaga pumilio*, which exhibits over 230 distinct alkaloids over its Central American range (Daly *et al*., 1987, 2002; Saporito *et al*., 2006, 2007a). First, we develop correlative models that approximate the spatial distribution of toxic ants, a crucial source of alkaloids for poison frogs (Saporito *et al*., 2012; Santos *et al*., 2016). Then, we apply Generalized Dissimilarity Modeling (GDM) (Ferrier *et al*., 2007) to assess the contribution of projected ant assemblage turnover to chemical trait dissimilarity among sites that have been screened for amphibian alkaloids, thus treating these toxins as a “community of traits”. To examine the effects of evolutionary relatedness, we implement a Multiple Matrix Regression approach (MMRR) (Wang, 2013) that incorporates not only prey turnover but also genetic divergences between *O. pumilio* populations as inferred from a neutral genetic marker.

## Material and Methods

### Estimating frog poison composition dissimilarity

As poison composition data of *Oophaga pumilio*, we used lipophilic alkaloid profiles derived from gas chromatography coupled with mass spectrometry of 934 frog skins sampled in 53 Central American sites (Daly *et al*., 1987, 2002; Saporito *et al*., 2006, 2007a), as compiled by Saporito *et al*. (2007a). We georeferenced sampled sites based on maps and localities presented by Saporito *et al*. (2007a) and other studies of *O. pumilio* (Saporito *et al*., 2006; Wang and Shaffer, 2008; Brown *et al*., 2010; Hauswaldt *et al*., 2011; Gehara *et al*., 2013). Because haphazard sampling may exaggerate poison composition variation in this dataset, and to maximize alkaloid sampling effort, we combined data from different expeditions to each site, pooling data from individuals. As such, we do not focus on potential short-term individual fluctuations in toxin composition (Saporito *et al*., 2006), but instead on variation tied to spatial gradients. To match the finest resolution available for the data grid predictors (see below), we combined alkaloid data within a 1 km^2^ grid cell. The final dataset included 230 alkaloids from 21 structural classes in 46 grid cell sites (Fig. 1) (alkaloid and locality data to be presented in the Supporting Information 1; presentation of raw data pending manuscript acceptance. See Supporting Information 2 for decisions on alkaloid identity). To estimate matrices of alkaloid composition dissimilarity (pairwise Sorensen’s distances) across frog populations, as well as geographic distances between sites, we used the *fossil* package (Vavrek, 2011) in R v. 3.3.3 (R Development Core Team, 2018).

**Figure 1.**
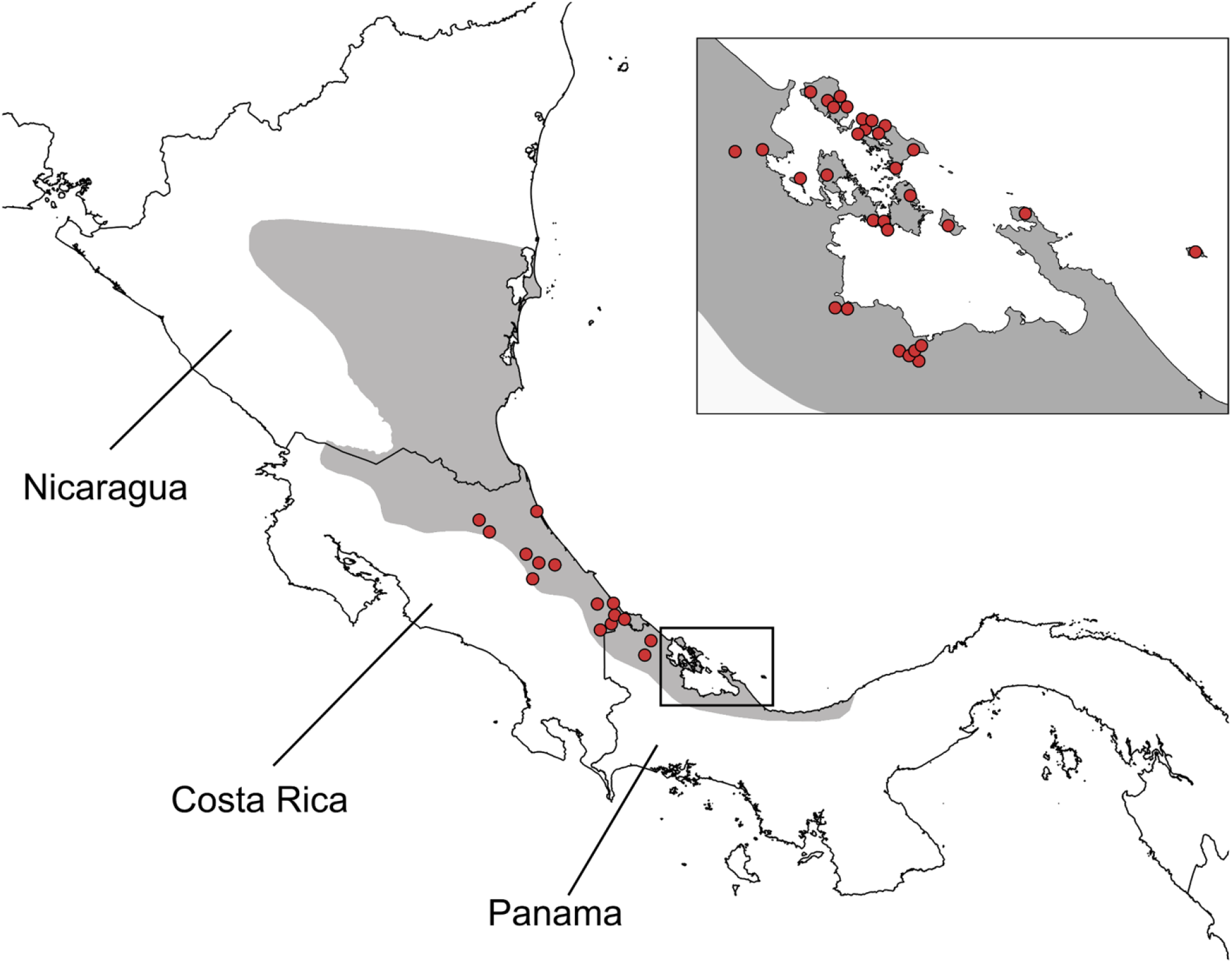
Sites sampled for skin alkaloid profiles of the strawberry poison frog, *Oophaga pumilio*. Each site corresponds to a 1 km^2^ grid cell, matching the resolution of environmental predictors. Original alkaloid data compiled by Saporito *et al*. (2017a). The distribution of *O. pumilio* is indicated in dark grey.

### Estimating arthropod prey assemblage dissimilarity

To approximate the spatial turnover of prey available to *Oophaga pumilio*, we focused on alkaloid-bearing ants that occur in (but are not necessarily restricted to) Costa Rica, Nicaragua and Panama, where that frog occurs. Ants are *O. pumilio*’s primary prey type, corresponding to more than half of the ingested volume (Donnelly, 1991; Caldwell, 1996; Darst *et al*., 2004). This frog also eats a large proportion of mites; however, limited occurrence data and taxonomic knowledge for mite species (e.g., McGugan *et al*., 2016) precluded their inclusion in our spatial analyses. Following a comprehensive literature search of alkaloid occurrence in ant taxa (Ritter *et al*., 1973; Wheeler, 1981; Jones *et al*., 1982a,b, 1988, 1996, 1999, 2007, 2012; Daly *et al*., 1994, 2000, 2005; Schroder *et al*., 1996; Spande *et al*., 1999; Leclercq *et al*., 2000; Saporito *et al*., 2004, 2009; Clark *et al*., 2005; Fox *et al*., 2012; Chen *et al*., 2013; Adams *et al*., 2015; Touchard *et al*., 2016), we estimated ant composition dissimilarity based on species that belong to genera known to harbor alkaloids, as follows: *Acromyrmex, Anochetus*, *Aphaenogaster*, *Atta, Brachymyrmex*, *Megalomyrmex*, *Monomorium*, *Nylanderia*, *Solenopsis*, and *Tetramorium* (see Supporting Information 2 for decisions on alkaloid presence in ant taxa). Georeferenced records were compiled from the Ant Web database as per June 2017 using the *antweb* R package (AntWeb, 2017). The search was restricted to the continental Americas between latitudes 40°N and 40°S. We retained only those 68 ant species that had a minimum of five unique occurrence records after spatial rarefaction (see below); the final dataset included a total of 1,417 unique records. Ant locality data to be presented in the Supporting Information 3 (presentation of raw data pending manuscript acceptance).

Because available ant records rarely matched the exact locations where *O. pumilio* alkaloids were characterized, we modeled the distribution of ant species at the landscape level to allow an estimation of prey composition at sites with empirical frog poison data. We created a species distribution model (SDM) for each ant species using MaxEnt (Phillips *et al*., 2006) based on 19 bioclimatic variables from the Worldclim v. 1.4 database (Hijmans *et al*., 2005; available at www.worldclim.org). Before modeling, we used the *spThin* R package (Aiello-Lammens, 2015) to rarefy ant records and ensure a minimum distance of 5 km between points, thus reducing environmental bias from spatial auto-correlation (Veloz, 2009; Boria *et al*., 2014). To reduce model over-fitting, we created a minimum-convex polygon defined by a 100 km radius around each species occurrence points, restricting background point selection by the modeling algorithm (Phillips *et al*., 2009; Anderson and Raza, 2010).

To properly parameterize the individual ant SDMs (Shcheglovitova and Anderson, 2013; Boria *et al*., 2014), we chose the best combination of feature class (shape of the function describing species occurrence vs. environmental predictors) and regularization multiplier (how closely a model fits known occurrence records) using the *ENMeval* R package (Muscarella *et al*., 2014). For species with 20 or more records, we evaluated models using k-fold cross-validation, which segregates training and testing points in different random bins (in this case, k = 5). For species with less than 20 records, we evaluated model fit using jackknife, a particular case of k-fold cross validation where the number of bins (k) is equal to the total number of points. We evaluated model fit under five combinations of feature classes, as follows: (1) Linear, (2) Linear and Quadratic, (3) Hinge, (4) Linear, Quadratic and Hinge, and (5) Linear, Quadratic, Hinge, Product, and Threshold. As regularization multipliers, we tested values ranging from 0.5 to 5 in 0.5 increments (Shcheglovitova and Anderson, 2013; Brown, 2014). For each model, the best parameter combination was selected using AICc. Based on this best combination, we generated a final SDM for each ant species (parameters used in all SDMs are presented in the Supporting Information 4).

To estimate a matrix of prey assemblage turnover based on ant distributions, we converted SDMs to binary maps using the 10^th^ percentile presence threshold. We then extracted presences for each ant species at sites sampled for frog alkaloids using the *raster* R package (Hijmans and Van Etten, 2016). Matrices of estimated ant composition dissimilarity (pairwise Sorensen’s distances) were calculated with *fossil* in R. To visualize ant species turnover across the range of *O. pumilio*, we reduced the final dissimilarity matrices to three ordination axes by applying multidimensional scaling using the *cmdscale* function in R. Each axis was then assigned a separate RGB color (red, green, or blue) as per Brown *et al*. (2014). A distribution layer for *O. pumilio* was obtained from the IUCN database (available at www.iucn.org).

### Modeling alkaloid composition turnover as a function of ant species turnover

To test the hypothesis that toxin composition in poison frogs vary geographically as a function of the composition of prey species, we modeled the turnover of *O. pumilio* alkaloids using a Generalized Dissimilarity Modeling (GDM) approach (Ferrier *et al*., 2007). GDM is an extension of matrix regression developed to model species composition turnover across regions as a function of environmental predictors. Once a GDM model is fitted to available biological data (estimated from species presences at sampled sites), the compositional dissimilarity across unsampled areas can be estimated based on environmental predictors (available for both sampled and unsampled sites). We implemented GDM following the steps of Rosauer *et al*. (2014) using the *GDM* R package (Manion *et al*., 2018).

We used the compiled database of alkaloids per site as the base of our GDMs. As predictor variables, we initially included geographic distance and all 68 individual models of ant species distributions at a 1 km^2^ resolution. To select the combination of predictors that contribute the most to GDM models while avoiding redundant variables, we implemented a stepwise backward elimination process (Williams *et al*., 2012), as follows: first, we built a model with all predictor variables; then, those variables that explained less than the arbitrary amount of 0.1% of the data deviance were removed iteratively, until only those variables contributing more than 0.1% were left.

To visualize estimated alkaloid turnover on geographic space from GDM outputs, we applied multidimensional scaling on the resulting dissimilarity matrices, following the procedure outlined above for the ant SDMs.

### Estimating frog population genetic divergence

To evaluate associations between alkaloid composition and shared evolutionary history between frog populations, we used the *cytochrome B* gene dataset of Hauswaldt *et al*. (2011), who sampled 197 *O. pumilio* individuals from 25 Central American localities. Because most sites sampled for genetic data are geographically close to sites sampled for alkaloids (Saporito *et al*., 2007a; Hauswaldt *et al*., 2011), we paired up alkaloid and genetic data based on Voronoi diagrams. A Voronoi diagram is a polygon whose boundaries encompass the area that is closest to a reference point relative to all other points of any other polygon (Aurenhammer, 1991). Specifically, we estimated polygons using the sites sampled for genetic data as references points and paired them with the sites for toxin data contained within each resulting Voronoi diagram. To estimate Voronoi diagrams, we used ArcGIS 10.3 (ESRI, Redlands). To calculate a matrix of average uncorrected pairwise genetic distances among localities, we used Mega 7 (Kumar *et al*., 2016). Six genetically sampled sites that had no corresponding alkaloid data were excluded from the analyses.

### Testing associations between toxin composition, prey assemblage dissimilarity, and population genetic divergence

To test the effect of population evolutionary divergence on poison composition of *O. pumilio*, we implemented a Multiple Matrix Regression with Randomization (MMRR) approach in R (Wang, 2013). To allow comparisons between genetic divergence and prey composition, we also included a matrix of ant assemblage dissimilarity as a predictor variable in MMRR models. As response variables, we focused on two distinct alkaloid datasets, namely individual alkaloids (n = 230) and alkaloid structural classes (n = 21). Additionally, to account for variation in poison composition resulting from ingestion of arthropod sources other than ants (i.e., mites, beetles, millipedes; Saporito *et al*., 2009), we performed a second set of analyses restricted to alkaloids (n = 125) belonging to alkaloid classes (n = 9) reported to occur in ant taxa. To assess the statistical significance of MMRR models and predictor variables, 10,000 permutations were used.

Matrices of alkaloid composition dissimilarity, estimated ant assemblage dissimilarity, genetic distances between *O. pumilio* populations, and geographic distances between sites, as well as Supporting Information and R scripts used in all analyses, are available online through GitHub (https://github.com/ivanprates/2019_gh_pumilio).

## Results

### Toxin diversity

Compilation of toxin profiles revealed large geographic variation in the number of alkaloids that compose the poison of *Oophaga pumilio*. The richness of individual alkaloids across sites varied between 7 and 48, while the richness of alkaloid structural classes varied between 3 and 17. Alkaloid composition turnover among sites was high; pairwise Sorensen distances ranged from 0.33 to 1 for individual alkaloids, and from 0.08 to 0.86 for alkaloid structural classes (Sorensen distances vary from 0 to 1, with 0 representing identical compositions and 1 representing no composition overlap across sites). A matrix regression framework using MMRR indicated that geographic distances affected the turnover of both individual alkaloids (p < 0.0001; R^2^ = 0.16) and alkaloid structural classes (p = 0.003; R^2^ = 0.09).

### Ant composition turnover

Distribution modeling of ant species suggested variation of ant richness over the landscape, with a concentration of species in the central portion of the distribution of *O. pumilio* in Costa Rica and southern Nicaragua (Fig. 2). The estimated number of alkaloid-bearing ant species at sites sampled for toxin profiles ranged between 24 and 52. However, ant richness did not have a significant effect on the number of individual alkaloids (linear regression; p > 0.22) (Fig. 3a) or alkaloid structural classes (p > 0.07).

**Figure 2.**
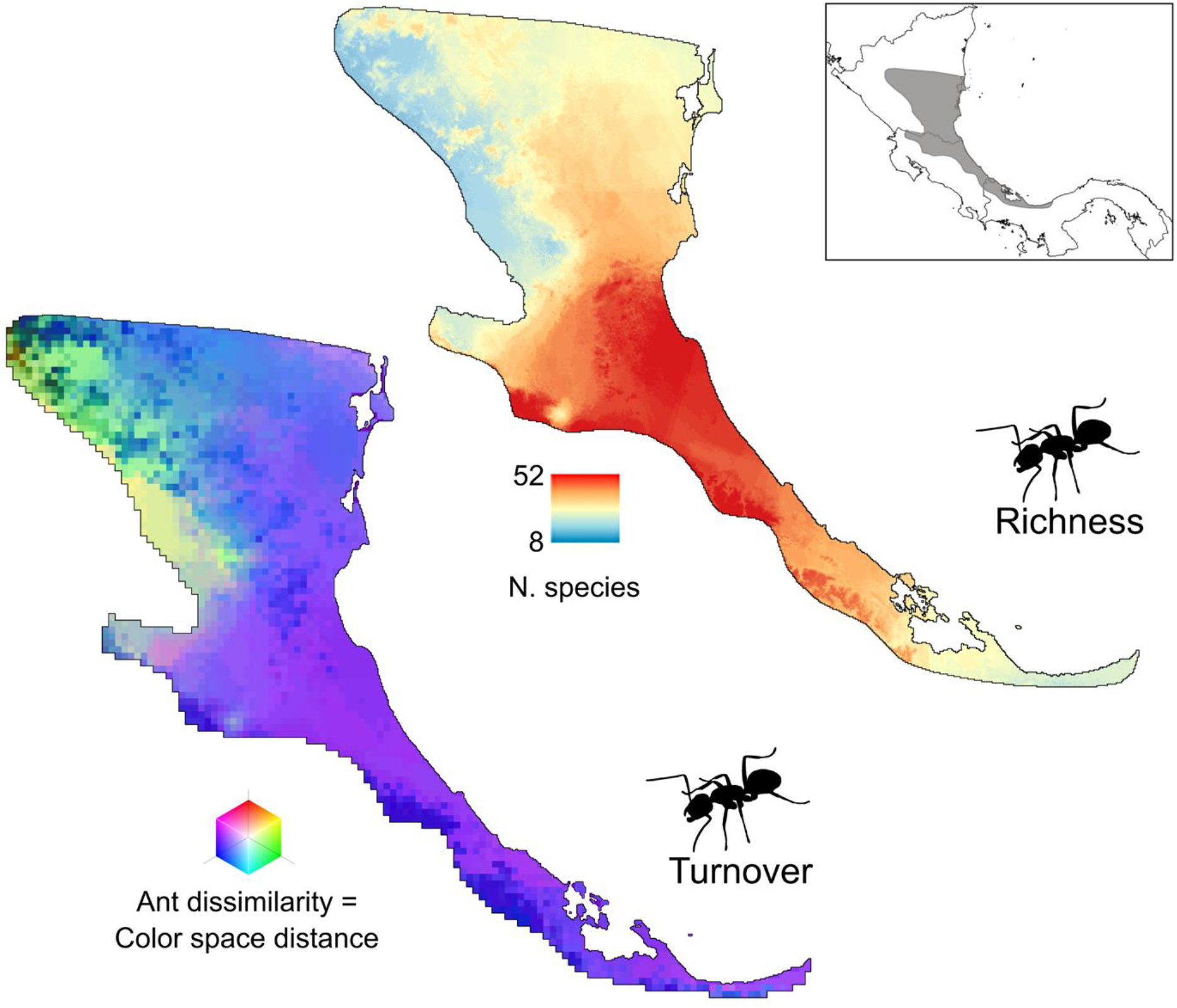
Estimated species turnover (left) and richness of prey assemblages across the range of the poison frog *Oophaga pumilio* based on projected distributions of 68 ant species from alkaloid-bearing genera. Inset indicates the natural range of *O. pumilio* (dark grey).

**Figure 3.**
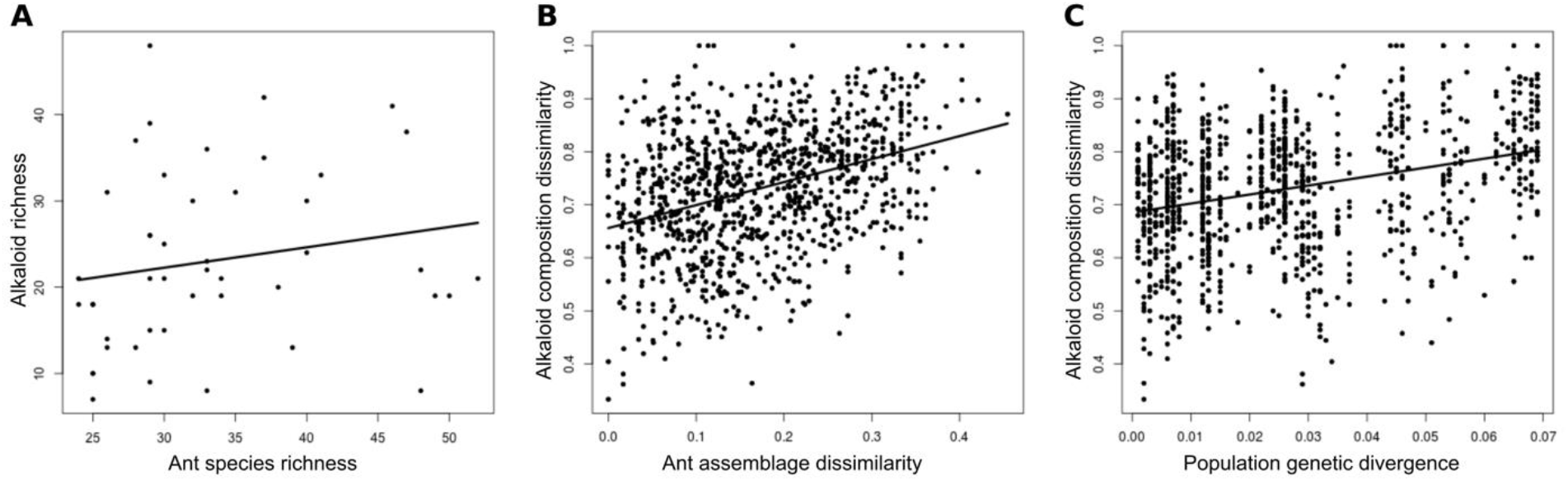
Alkaloid richness in *Oophaga pumilio* as a function of ant assemblage richness across sites (A); alkaloid dissimilarity in *O. pumilio* as a function of ant assemblage dissimilarity across sites (B); and alkaloid dissimilarity in *O. pumilio* as a function of population genetic divergence based on a neutral marker (C). Relationships in B and C are statistically significant; significance was estimated from a Multiple Matrix Regression with Randomization (MMRR) approach (see text).

Estimated ant assemblage turnover was low within the distribution of *O. pumilio* (Fig. 2). Pairwise ant Sorensen distances ranged from 0 to 0.47 at sites sampled for frog alkaloids, reflecting the broad ranges inferred for several alkaloid-bearing ant species. MMRR analyses suggested a significant effect of geographic distances between sites on species turnover of alkaloid-bearing ants (p < 0.0001; R^2^ = 0.62).

Projection of ant beta-diversity on geographic space suggested that prey assemblages are similar throughout the southern range of *O. pumilio* in Panama and Costa Rica, with a transition in the northern part of the range in inland Nicaragua (Fig. 2). Inner mid-elevations are expected to harbor ant assemblages that are distinct from those in the coastal lowlands.

### Generalized Dissimilarity Modeling

Dissimilarity modeling supports the idea that ant assemblage turnover affects the spatial variation of poison composition in *O. pumilio*. A GDM model explained 22.9% of the alkaloid composition turnover when the model included geographic distances among sampled sites. A model that did not incorporate geographic distances explained 20% of alkaloid profile dissimilarity. After eliminating those ant distribution models that contributed very little to the GDM (< 0.1%) using a stepwise backward elimination procedure, only nine out of 68 species were retained in the final model, as follows: *Acromyrmex coronatus*, *Anochetus orchidicola*, *Brachymyrmex coactus*, *Brachymyrmex pictus*, *Monomorium ebeninum*, *Solenopsis azteca*, *Solenopsis bicolor*, *Solenopsis pollux*, and an undescribed *Solenopsis* species (sp. “jtl001”). A GDM model not including geographic distances explained 20% of the observed alkaloid beta-diversity also for this reduced ant dataset.

Projection of GDM outputs onto geographic space indicates areas expected to have more similar amphibian alkaloid profiles based on ant turnover (Fig. 4). The results suggest latitudinal variation in poison composition, with a gradual transition from the southern part of the distribution of *O. pumilio* (in Panama) through the central and northern parts of the range (in Costa Rica and Nicaragua, respectively) (pink to purple to blue in Fig. 4). Another major transition in poison composition was estimated across the eastern and western parts of the range of *O. pumilio* in Nicaragua (blue to green in Fig. 4).

**Figure 4.**
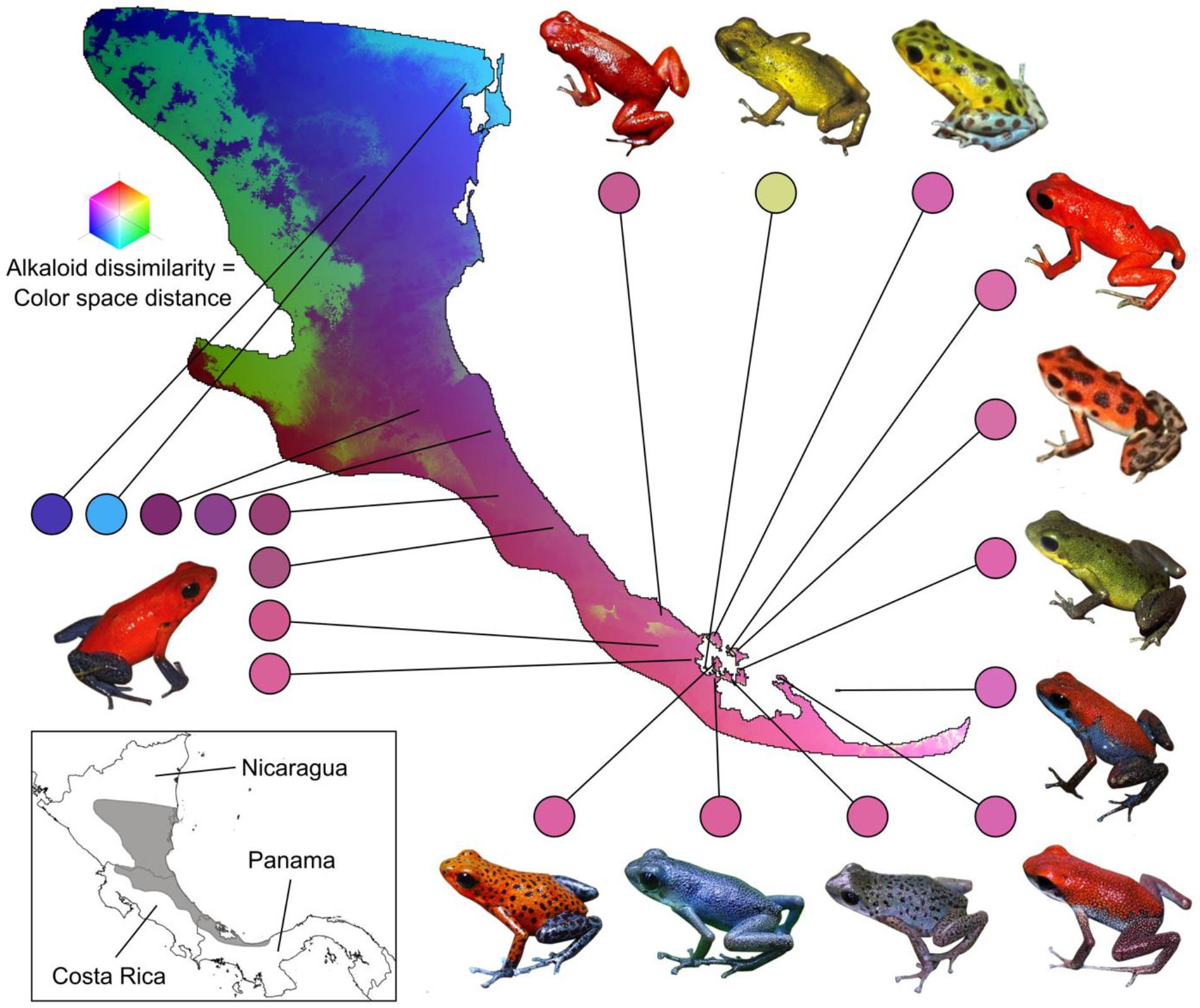
Estimated composition dissimilarity of alkaloid profiles across the range of *Oophaga pumilio* as a function of spatial turnover of alkaloid-bearing ant species, from a Generalized Dissimilarity Modeling approach (GDM). Similar colors on the map indicate similar estimated alkaloid profiles. Pictures indicate dorsal skin coloration patterns in *O. pumilio*. Inset indicates the natural range of *O. pumilio* (dark grey).

### Multiple Matrix Regressions

In agreement with the GDM results, MMRR analyses supported the hypothesis that spatial turnover of *O. pumilio* toxin profiles is affected by prey composition dissimilarity, in spite of the limited variation of ant assemblages. The estimated turnover of alkaloid-bearing ant assemblages had a significant effect on individual alkaloid composition dissimilarity (p < 0.0001; R^2^ = 0.13) (Fig. 3b), a result that changed little when considering only those alkaloids from structural classes known to occur in ants (p < 0.0001; R^2^ = 0.11). Similarly, estimated alkaloid-bearing ant assemblages had a significant yet weak effect on the dissimilarity of alkaloid structural classes among sites (p = 0.002; R^2^ = 0.05), a result that held when considering only classes known to occur in ants (p = 0.007; R^2^ = 0.03).

Genetic distances between populations of *O. pumilio* had a weak yet significant effect on frog alkaloid composition dissimilarity (p < 0.0001; R^2^ = 0.09) (Fig. 3c). A MMRR model incorporating both frog population genetic distances and ant composition dissimilarity explained around 15% of alkaloid composition variation among populations of *O. pumilio* (p_ants_ < 0.001; p_gen_ = 0.05).

## Discussion

### Effects of prey turnover on trait variation and its ecological consequences

Based on estimates of the distribution of alkaloid-bearing ants, we found associations between prey composition gradients and spatial turnover in the defensive chemical traits of the strawberry poison frog, *Oophaga pumilio*. Species distribution modeling supported that the pool of alkaloid sources varies in space. Moreover, GDM and MMRR analyses inferred that alkaloid beta diversity in *O. pumilio* covaries with the composition of ant assemblages. These results are consistent with observations that distinct toxins are restricted to specific arthropod taxa (Saporito *et al*., 2007a, 2012); prey species with limited distributions may contribute to unique defensive phenotypes in different parts of the range of poison frog species. Accordingly, GDM predicted unique alkaloid combinations in the northern range of *O. pumilio* (Nicaragua) relative to central and southern areas (Costa Rica and Panama) (Fig. 4); that northern region remains poorly known in terms of chemical diversity (Mebs *et al*., 2008), and future surveys may reveal novel toxin combinations in the poison frogs that occur therein. These results are consistent with the view that biotic interactions play a role in phenotypic variation within species and across space and influence functional diversity in ecological communities (Miner *et al*., 2005; Pelletier *et al*., 2009; Post and Palkovack, 2009).

Our analyses also indicate that different prey species may have disparate contributions to observed chemical trait variation in poison frogs. After eliminating those ants that contributed very little to the GDM (< 0.1%), only nine out of 68 species were retained in the final model. This result may imply that a few prey items contribute disproportionately to the uniqueness of chemical defenses among populations of *O. pumilio*. Alternatively, it may point to redundancy in the alkaloids provided by prey species to poison frogs. Specifically, it is possible that some of the ant species contributed less to alkaloid beta-diversity when in the presence of a co-distributed and potentially functionally equivalent species, therefore being removed during the GDM’s stepwise backward elimination procedure.

These results have potential implications for the ecology and evolution of the interacting species. For instance, frogs may favor and seek prey types that provide unique chemicals or chemical combinations, increasing survival rates from encounters with their predators. This idea is consistent with evidence that alkaloid quantity, type, and richness result in differences in the perceived palatability of poison frogs to predators (Bolton *et al*., 2017; Murray *et al*., 2017). Moreover, because alkaloids vary from mildly unpalatable to lethally toxic (Daly *et al*., 2005; Santos *et al*., 2016), the composition of amphibian poisons may affect the survival and behavior of their predators following an attack (Darst and Cummings, 2006). From the perspective of arthropod prey, predator feeding preferences and foraging behavior may affect population densities and dynamics, potentially in a predator density-dependent fashion (Pelletier *et al*., 2009; Bolnick *et al*., 2011). Finally, functional redundancy among prey types may confer resilience to the defensive phenotypes of poison frogs, ensuring continued protection from predators despite fluctuations in prey availability. Future investigations of these topics will advance our understanding of how the phenotypic outcomes of species interactions affect ecological processes.

### Evolutionary relatedness and phenotypic similarity

In addition to the contribution of prey assemblages, the results support that phenotypic similarity between populations also varies with evolutionary relatedness. MMRR analyses indicated that population genetic divergence has a significant effect on alkaloid beta diversity in *O. pumilio*, an effect that may reflect physiological or behavioral differences. For instance, feeding experiments support that poison frog species differ in their capacity for lipophilic alkaloid sequestration and that at least a few species can perform metabolic modification of ingested alkaloids (Daly *et al*., 2003, 2005, 2009). Additionally, an effect of evolutionary divergence on toxin profiles may stem from population differences in foraging strategies or behavioral prey choice (Daly *et al*., 2000; Saporito *et al*., 2007a). Importantly, however, MMRR suggested that genetic divergence accounts for less variation in the defensive phenotypes of *O. pumilio* than alkaloid-bearing prey.

We employed genetic distances from a neutral marker as a proxy for shared evolutionary history, and do not imply that this particular locus contributes to variation in frog physiology. An assessment of how genetic variants influence chemical trait diversity in poison frogs will rely on sampling of functional genomic variation, for which the development of genomic resources (e.g., Rogers *et al*., 2018) will be essential. Nevertheless, our approach can be extended to other systems where information on both neutral and functional genetic variation is available. These systems include, for instance, predators where nucleotide substitutions in ion channel genes confer resistance to toxic prey at a local scale (Feldman *et al*., 2010), and species where variation in major histocompatibility complex loci drives resistance to pathogens locally (Savage and Zamudio, 2016).

### Other sources of phenotypic variation

Although we show that prey community composition explains part of the spatial variation in frog poisons, this proportion is relatively small. This result may stem from challenges in quantifying turnover of prey species that act as alkaloid sources. Due to restricted taxonomic and distribution information, we estimated species distribution models only for a set of well-sampled ant species and were unable to include other critical sources of dietary alkaloids, particularly mites (Saporito *et al*., 2012; McGugan *et al*., 2016). Although not being able to incorporate a substantial fraction of the prey assemblages expected to influence poison composition in *O. pumilio*, the GDM was able to explain about 23% of poison variation in this species. This proportion is likely to increase with the inclusion of additional species of ants and other alkaloid-bearing prey taxa, such as mites, beetles, and millipedes (Daly *et al*., 2000; Saporito *et al*., 2003; Dumbacher *et al*., 2004).

Limitations in our knowledge of arthropod chemical diversity may also have affected the analyses. Contrary to our expectations, we found a slight decrease in the explanatory value of models that incorporated only alkaloids from structural classes currently known to occur in ants, a result that may reflect an underestimation of ant chemical diversity. For instance, mites and beetles are thought to be the source of tricyclics to Neotropical poison frogs, but the discovery of these alkaloids in African Myrmicinae ants suggests that they may also occur in ants from other regions (Schroder *et al*., 1996). As studies keep describing naturally occurring alkaloids, we are far from a complete picture of chemical trait diversity in both arthropods and amphibians (Saporito *et al*., 2012).

This study demonstrates that incorporating species interactions can provide new insights into the drivers of phenotypic diversity even when other potential sources of variation are not fully understood. It is possible that frog poison composition responds not only to prey availability but also to alkaloid profiles in prey, since toxins may vary within arthropod species (Daly *et al*., 2002; Saporito *et al*., 2004, 2007b; Dall’Aglio-Holvorcem *et al*., 2009; Fox *et al*., 2012). There is evidence that arthropods synthesize their alkaloids endogenously (Leclercq *et al*., 1996; Camarano *et al*., 2012; Haulotte *et al*., 2012), but they might also sequester toxins from plants, fungi, and symbiotic microorganisms (Saporito *et al*., 2012; Santos *et al*., 2016). Accounting for these other causes of variation will be a challenging task to chemical ecologists. An alternative to bypass knowledge gaps on the taxonomy, distribution, and physiology of these potential toxin sources may be to focus on their environmental correlates. A possible extension of our approach is to develop models that incorporate abiotic predictors such as climate and soil variation, similarly to investigations of community composition turnover at the level of species (Ferrier *et al*., 2007; Zamborlini-Saiter *et al*., 2016).

### Eco-evolutionary feedbacks

Associations between prey assemblage composition and predator phenotype provide opportunities for eco-evolutionary feedbacks, the bidirectional effects between ecological and evolutionary processes (Post and Palkovacs, 2009). In the case of poison frogs, there is evidence that toxicity correlates to the conspicuousness of skin color patterns (Maan and Cummings, 2011), which therefore act as aposematic signals for predators (Hegna *et al*., 2013). However, coloration patterns also mediate assortative mating in poison frogs (Maan and Cummings, 2009; Crothers and Cummings, 2013). When toxicity decreases, the intensity of selection for aposematism also decreases, and mate choice may become a more important driver of color pattern evolution than predation (Summers *et al*., 1999). Divergence due to sexual selection may happen quickly when effective population sizes are small and population gene flow is limited, which is the case in *O. pumilio* (Gehara *et al*., 2013). By affecting chemical defenses, it may be that spatially structured prey assemblages have contributed to the vast diversity of color patterns seen in poison frogs. On the other hand, our GDM approach inferred similar alkaloid profiles between populations of *O. pumilio* that have distinct coloration patterns (Fig. 4). However, it may be challenging to predict frog toxicity from chemical profiles (Daly *et al*., 2005), and future studies of this topic will advance our understanding of the evolutionary consequences of poison composition variation.

### Concluding remarks

Integrative approaches have demonstrated how strong associations between species can lead to tight covariation among species traits, with an iconic example being that of predator and prey species that engage in “coevolutionary arms races” (Thompson, 2005). However, it may be harder to assess the outcomes of weaker interactions among multiple co-distributed organisms (Anderson, 2017). The chemical defense system of poison frogs varies as a function of prey assemblages composed of tens of arthropod species, each of them providing single or a few toxin molecules (Daly *et al*., 2005). Not surprisingly, it has been hard to predict the combinations of traits emerging from these interactions (Santos *et al*., 2016). The landscape ecology framework presented here approaches this problem by incorporating data on biotic interactions throughout the range of a focal species, a strategy that has also improved correlative models of species distributions (Lewis *et al*., 2017; Sanín and Anderson, 2018). Our framework can be extended to a range of systems that have similar structures, including other multi-trophic interactions (Van der Putten *et al*., 2001; Del-Claro, 2004; Scherber *et al*., 2010), geographic coevolutionary mosaics (Greene and McDiarmid, 1981; Mallet *et al*., 1995; Symula *et al*., 2001), microbial assemblages (Zomorrodi and Maranas, 2012; Landesman *et al*., 2014), and ecosystem services (Moorhead and Sinsabaugh, 2006; Allison, 2012; Gossner *et al*., 2016). Integrative approaches to the problem of how spatially structured biotic interactions contribute to phenotypic diversity will continue to advance our understanding of the interplay between ecological and evolutionary processes.

## Acknowledgments

We are grateful to entomologists David Lohman, Corrie Moreau, and Israel del Toro for suggestions and advice about ant diversity, as well as its ecological drivers, in the initial stages of this project. Suggestions by Rayna C. Bell, Kevin P. Mulder, Michael L. Yuan, Edward A. Myers, Ryan K. Schott, Maria Strangas, Danielle Rivera, Catherine Graham, Robert Anderson, and the New York Species Distribution Modeling Group greatly improved this manuscript. This work was co-funded by FAPESP (BIOTA 2013/50297–0), NSF (DEB 1343578), and NASA through the Dimensions of Biodiversity Program. IP acknowledges additional funding from a Smithsonian Peter Buck Postdoctoral Fellowship.

## Authorship statement

AC acquired funding and supervised the research team. IP, AP, and JLB designed and performed the analyses, wrote software, and worked on the visualization of results. IP obtained and curated the data and led the writing of the original draft. All four authors conceptualized the study, interpreted the results, and contributed to manuscript writing.

## Data statement

All raw data, dissimilarity matrices, and Supporting Information will be made available online through both Dryad and GitHub (https://github.com/ivanprates/2019_gh_pumilio). R scripts used in all analyses are provided in GitHub.

## Supporting Information

**Supporting Information 1.** Raw alkaloid and locality data [presentation of raw data pending manuscript acceptance].

**Supporting Information 2.** Decisions on alkaloid identity and alkaloid presence in ant taxa.

**Supporting Information 3.** Raw ant locality data used in species distribution models [presentation of raw data pending manuscript acceptance].

**Supporting Information 4.** Parameters used in all final ant species distribution models following optimization.

